# Structural and functional features characterizing the brains of individuals with higher controllability of motor imagery

**DOI:** 10.1101/2023.10.11.560970

**Authors:** Tomoya Furuta, Tomoyo Morita, Gen Miura, Eiichi Naito

## Abstract

Motor imagery is a higher-order cognitive brain function that mentally simulates movements without performing the actual physical one. Although many studies have dealt with motor imagery, neural bases that determine individual differences in motor imagery ability are not well understood. Using magnetic resonance imaging and controllability of motor imagery (CMI) test that can objectively evaluate individual ability to manipulate one’s imaginary postures, we show that the bilateral superior frontoparietal white matter region expands in higher CMI test scorers. In addition, CMI test activates the bilateral dorsal premotor cortex (PMD) and superior parietal lobule (SPL); specifically, the left PMD and/or the right SPL enhance functional coupling with the visual body, somatosensory, and motor/kinesthetic areas in the higher scorers. This study elucidated the structural and functional features that characterize the brains of individuals with higher controllability of motor imagery, and advanced understanding of individual differences in motor imagery ability.

## Introduction

Motor imagery is a higher-order cognitive brain function that internally simulates movements without performing the actual physical movements^1–5^. Motor imagery has attracted the interest of many psychologists and neuroscientists^6,7^, and mental practice utilizing motor imagery has been widely used in sports training and post-stroke rehabilitation^8–11^.

When applying motor imagery to mental practice, individual differences in motor imagery ability must be considered. As objective assessment of an individual’s motor imagery ability is difficult, in many previous studies, subjective measures of imagery vividness and easiness, e.g., the vividness of movement imagery questionnaire (VMIQ)^12^, movement imagery questionnaire (MIQ)^13^, or their revised versions (e.g., VMIQ-2^14^, revised MIQ-R^15^, or MIQ-RS^16^), are commonly used to assess individual ability of motor imagery. However, these are subjective assessments of the quality of an individual’s motor imagery and are not always objective assessments of an individual’s motor imagery ability, making it difficult to objectively investigate neural bases that determine individual differences in motor imagery ability. Recently, a more objective evaluation method was proposed^17^, and individual differences in motor imagery are gradually being unveiled^18,19^. Nevertheless, the structural and functional characteristics of the brain that determine individual differences in motor imagery ability are not well understood.

An individual’s ability to accurately manipulate and control motor imagery, i.e., controllability of motor imagery, is essential for motor imagery function. In this study, we introduce controllability of motor imagery (CMI) test^2,20–22^, which can objectively evaluate individual’s ability to accurately manipulate motor imagery. By using the CMI test, an individual’s ability to internally generate, manipulate, and hold one’s imaginary body postures from a first-person perspective in response to each of five consecutive verbal instructions regarding movements of body parts (left or right arms, left or right legs, upper body, and head/neck) can be evaluated. After the final instruction, participants were asked to perform the final posture by themselves. By evaluating the final posture, whether the participants could manipulate their motor imagery appropriately during the test can be assessed (Figure 1a). Unlike the subjective assessments of whether a participant can vividly or easily imagine a certain movement (see above), the CMI test provides a unique and reliable measure to objectively evaluate an individual’s ability to precisely manipulate one’s own postural image, i.e., motor imagery. Thus, this study aimed to elucidate structural and functional features that characterize the brains of individuals with higher controllability of motor imagery, using the CMI test and magnetic resonance imaging (MRI).

**Figure 1.**
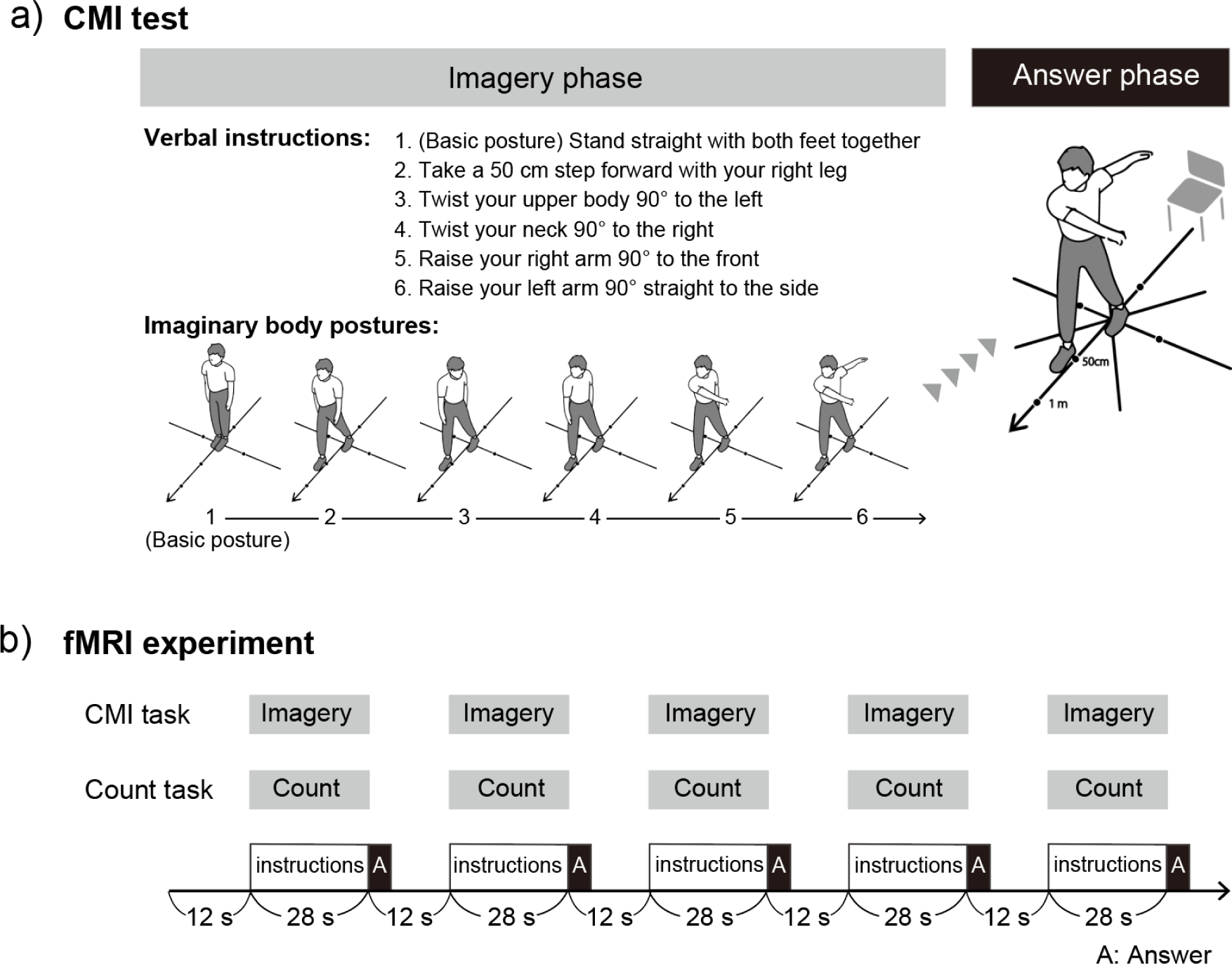
a) An example of a trial of CMI test and b) task design in fMRI experiment. Abbreviations: A, Answer phase; CMI, controllability of motor imagery.

A meta-analysis of functional neuroimaging studies for motor imagery elucidated the consistent importance of the frontoparietal and subcortical motor-related brain regions during various types of motor imagery^6^. Among these brain regions, motor imagery function is severely impaired in patients with frontal (premotor) and parietal injuries^23–25;^ however, it is preserved without these damages^26^. These lines of evidence indicate the importance of frontoparietal regions that are most likely connected by the three branches of the superior longitudinal fasciculus (SLF I, II, and III for superior, middle, and inferior branches, respectively)^27–29^ in motor imagery.

Among frontoparietal regions, many studies have consistently reported activations in the dorsal premotor cortex (PMD) and the superior parietal lobule (SPL) during motor imagery^4,29–36^. The PMD and SPL forms a functional network connected by the SLF I or II^27,29^. Furthermore, among the SLF (I, II, and III), the preferential role of the brain regions connected by SLF I or II for spatial/motor processing, particularly the greater contribution of the regions connected by SLF I for mental motor imagery, was demonstrated^29^.

Based on this knowledge, we first test our hypothesis whether the brain structures (white and gray matter) associated with SLF I or II expands in higher scorers on the CMI test (=participants with higher controllability of motor imagery). In addition, if the PMD–SPL network plays central roles for controllability of motor imagery, we may expect that a component of mental manipulation of motor imagery during the CMI test activates the PMD–SPL network, and the higher scorers on the CMI test show greater imagery-related activity in this network. Finally, we also investigated possible brain regions that enhanced functional coupling with this network in higher scorers. We address these points by collecting and analyzing T1-weighted structural and functional MRI data.

## Results

In this study, 89 healthy right-handed young adults performed the CMI test outside the MRI scanner room. The CMI test consists of 10 trials, each of which consists of five consecutive verbal instructions regarding movements of the six body parts (see Methods). Starting from the basic posture, participants with eyes closed imagined to move only one body part per instruction and kept updating their imaginary postures according to all instructions through a trial. The participants must performed the final posture, and experimenters evaluated its correctness. The average number of correct answers across 10 trials was calculated per participant (= CMI test score; maximum score = 6.0).

Moreover, 28 of the 89 participants performed the CMI task while brain activity was scanned. In this case, demonstrating the final posture inside the scanner was impossible, so they were asked to rate how confident they were about the accuracy of their imaginary final posture (subjective rating of confidence) by pressing a corresponding button after each trial.

The mean CMI test score outside the scanner room for the 89 participants was 4.6 ± 0.7 (range 2.7–6.0), and the mean score for the 28 participants assigned for the functional MRI experiment was 4.4 ± 0.8 (range 2.7–6.0). The correlation between the CMI test score and the subjective rating of confidence across 28 participants was not significant (r = 0.14, N = 28, p = 0.47). Thus, the subjective rating of confidence was not directly related to individual’s controllability of motor imagery objectively evaluated by the CMI test.

### White matter expansion in higher scorers on the CMI test

In 89 participants, to examine white and gray matter structures that expand in higher scorers on the CMI test, voxel-based morphometry (VBM) analysis was performed in the whole brain. The brain regions in which the tissue volume correlated with the score of the CMI test performed outside the MRI scanner room were identified.

In the white matter regions, we found a significant cluster of voxels that positively correlated with the score in each of the left and right hemispheres (Figure 2a– 2c). These were the only significant clusters in the entire brain and located in the white matter regions in which the fiber tract connecting the frontoparietal regions is likely running. Using publicly available probability maps of the tract (see Methods), we checked whether these clusters are located in the regions where the SLF (I, II, or III) is likely running. Of the total volume (12191 mm^3^) of the two clusters, 8972 mm^3^ (approximately 74%) overlapped with the white matter regions where the SLF (I, II, or III) is likely running (Figure 2d). Moreover, 28% (2512 mm^3^) of the 8972 mm^3^ was found to be in the regions where SLF I (including overlap with the SLF II or III) is likely running. Similarly, 83% (7440 mm^3^) of the 8972 mm^3^ was found to be in the regions where SLF II (including overlap with SLF I or III) is likely running. The majority (7835 mm^3^, 87%) of the 8972 mm^3^ was found to be in the regions where SLF I or II is likely running. In addition, 41% (3659 mm^3^) of the 8972 mm^3^ was found to be in the regions where SLF III (including overlap with the SLF I or II) is likely running.

**Figure 2.**
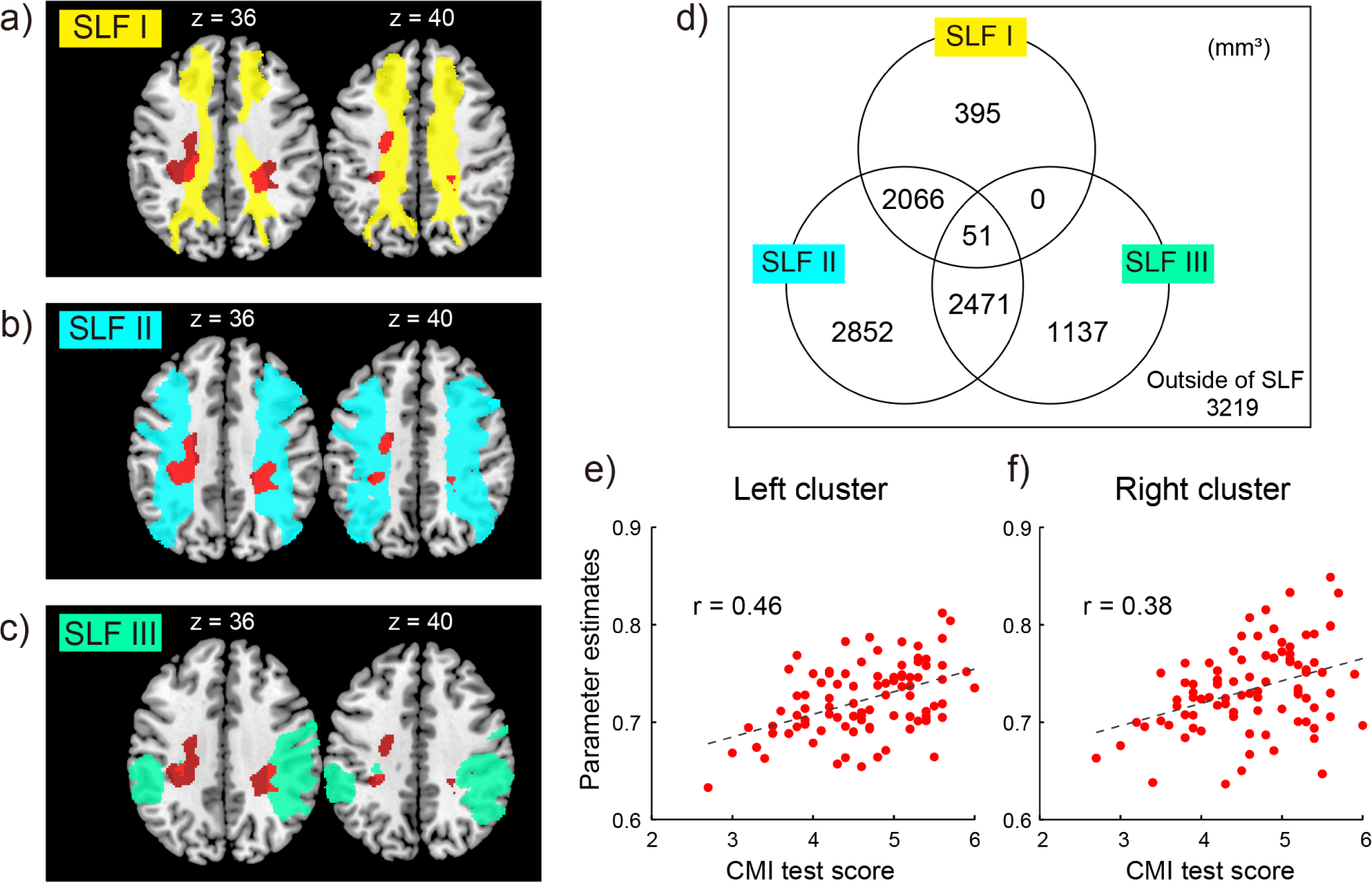
Results of the VBM analysis. **a–c)** Bilateral white matter clusters (red sections) whose volume positively correlated with the CMI test score performed outside of the MRI scanner room across 89 participants. The clusters and tract probability maps of SLF I (yellow sections; panel a), II (light blue sections; panel b), or III (light green sections; panel c) are superimposed on the horizontal sections (z = 36 and 40) of the MNI standard brain. **d)** Overlapping volume (mm^3^) between the white matter clusters and tract probability maps of the SLF (I, II, or III). **e, f)** Interindividual correlation between the normalized size (parameter estimates) of the left (panel e) or right (panel f) cluster (vertical axis) and the CMI test score (horizontal axis). Each dot represents individual data. Abbreviations: MNI, Montreal Neurological Institute; SLF, superior longitudinal fasciculus.

Then, the interindividual correlation between the normalized size of the left or right cluster and the CMI test score was visualized. A positive correlation was confirmed in each cluster (correlation coefficient r = 0.46, left cluster; r = 0.38, right cluster; Figure 2e, 2f). Viewed collectively, individuals with higher CMI test scores showed bilateral expansion of white matter regions where the SLF (I, II, or III) is most likely running.

We found that neither gray matter regions showed a positive correlation with the score nor white matter regions showed a negative correlation with the score.

### Brain regions active during the CMI task when compared with a control task

Of the 89 participants, 28 also performed the CMI task and a control (count) task while scanning brain activity (Figure 1b). In the count task, they listened to the same verbal instructions but kept counting (remembered) the number of term that “right” (right arm, right leg, and rightward) appeared in the instructions without imagining the postural changes. Thus, in this control task, the participants must pay attention to and process the same verbal instructions; however, its associated mental process was different from the mental imagery required during the CMI task. Thus, by comparing the brain activity during the CMI task with that during the count task, one may identify brain activity associated with mental imagery (manipulation) processes during the task (=imagery-related activity).

When we examined brain regions where activity increased during the CMI task compared with the count task, we found five significant clusters mainly in the bilateral superior frontoparietal cortices (Figure 3a–3c). Two of them were located in the bilateral PMD. The left parietal cluster located in the SPL was also extending to the postcentral gyrus, and the right-parietal cluster located in the SPL was extending to the intraparietal sulcus (IPS) and inferior parietal lobule (IPL). A cluster in the left lateral occipital cortex (LOC) was also found extending dorsally to the IPS and medially to the precuneus.

**Figure 3.**
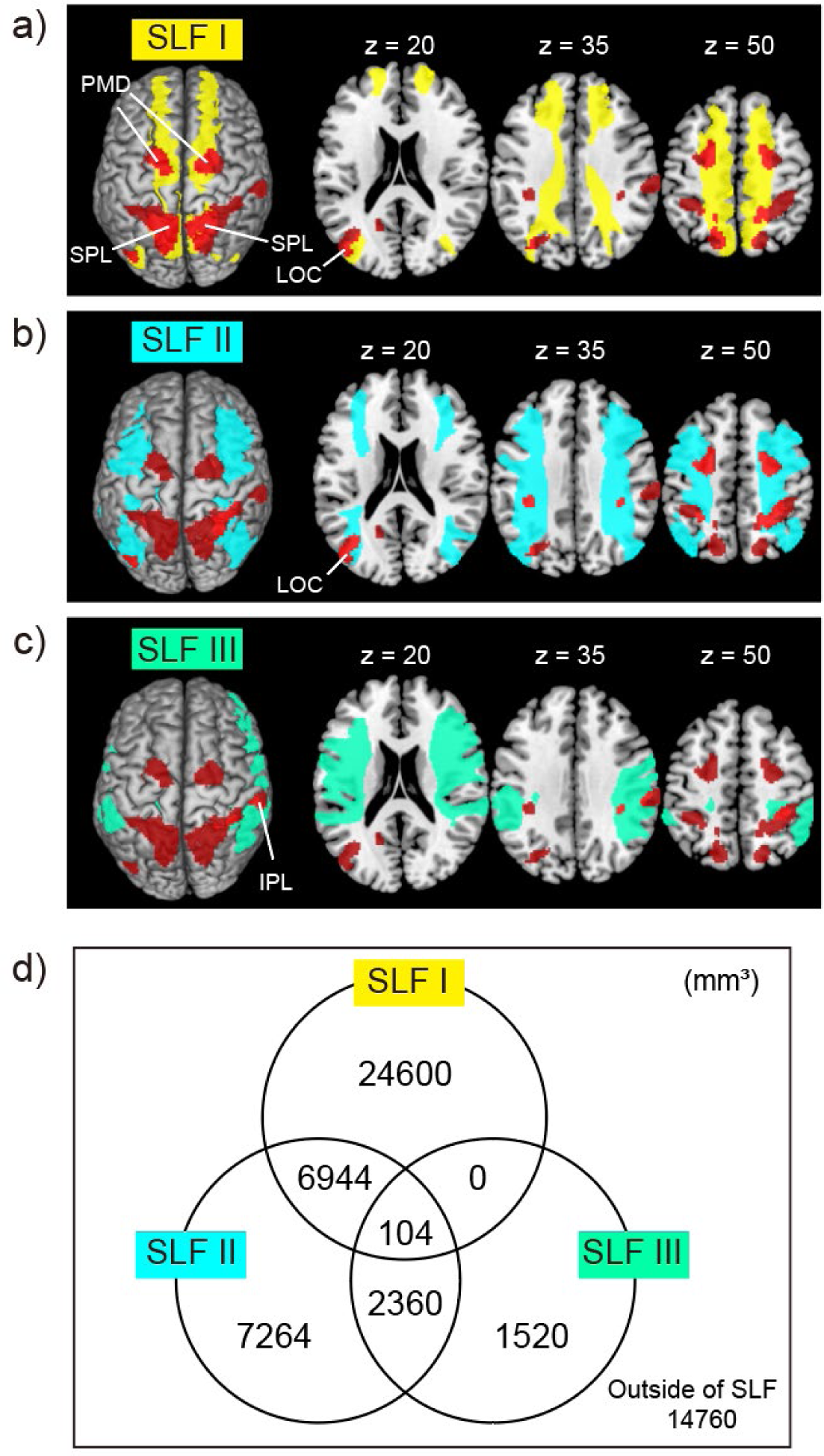
Imagery-related activity. **a–c)** The imagery-related activity (CMI task > count task; red sections) is superimposed on the MNI standard brain and its horizontal sections (z = 20, 35, and 50). The tract probability maps of SLF I (yellow sections; panel a), II (light blue sections; panel b), or III (light green sections; panel c) are also superimposed. **d)** Overlapping volume (mm^3^) between the imagery-related activity and tract probability maps of the SLF (I, II, or III). Abbreviations: IPL, inferior parietal lobule; LOC, lateral occipital cortex; MNI, Montreal Neurological Institute; PMD, dorsal premotor cortex; SLF, superior longitudinal fasciculus; SPL, superior parietal lobule.

Then, we checked whether these clusters are located in the regions likely connected by the SLF (I, II, or III), using the same probability maps described above (Figure 3d). Of the total volume (57552 mm^3^) of the five clusters, 42792 mm^3^ (approximately 74%) overlapped with the brain regions that are likely connected by the SLF (I, II, or III). In addition, 74% (31648 mm^3^) of the 42792 mm^3^ was found in the regions that are likely connected by SLF I (including overlaps with SLF II or III). Similarly, 39% (16672 mm^3^) of the 42792 mm^3^ was found in regions that are likely connected by SLF II (including overlaps with SLF I or III). Nearly all of 42792 mm^3^ (41272 mm^3^, 96%) was found in regions that are likely connected by SLF I or II. Finally, 9% (3984 mm^3^) of 42792 mm^3^ was found in regions that are likely connected by SLF III (including overlaps with SLF I or II). As we hypothesized, among the brain regions likely connected by the SLF (I, II, or III), the imagery-related activity was mainly located in the superior frontoparietal (PMD–SPL) regions likely connected by SLF I or II.

As in the VBM analysis, brain regions in which imagery-related activity correlated with the CMI test score were examined across 28 participants. However, no such regions were found.

### Functional connectivity

By conducting a generalized psychophysiological interaction analysis, we examined brain regions in which imagery-related functional coupling with a seed region changes in relation to the CMI test score across 28 participants. We prepared eight seed regions in the left PMD (cytoarchitectonic areas 6d3 and 6d1)^37^, right PMD (areas 6d3 and 6d1), left SPL (areas 5L and 7A), and right SPL (areas 5L and 7A) in bilateral PMD and SPL clusters identified above (Figure 3a–3c).

When the left area 6d3 was a seed region (Figure 4a), two significant clusters were found in the left-early visual cortices and higher-order visual cortices (extrastriate body area [EBA]) in which imagery-related connectivity positively correlated with the CMI test score (Figure 4b), whereas no significant clusters were found when the left area 6d1 was a seed region. For each cluster, the relationship between the CMI test score and connectivity was plotted for all participants, and a correlation coefficient of r = 0.58 was found in the early visual cluster and r = 0.69 in the EBA cluster (Figure 4c, 4d). When right areas 6d3 and 6d1 were seed regions, no significant clusters were found.

**Figure 4.**
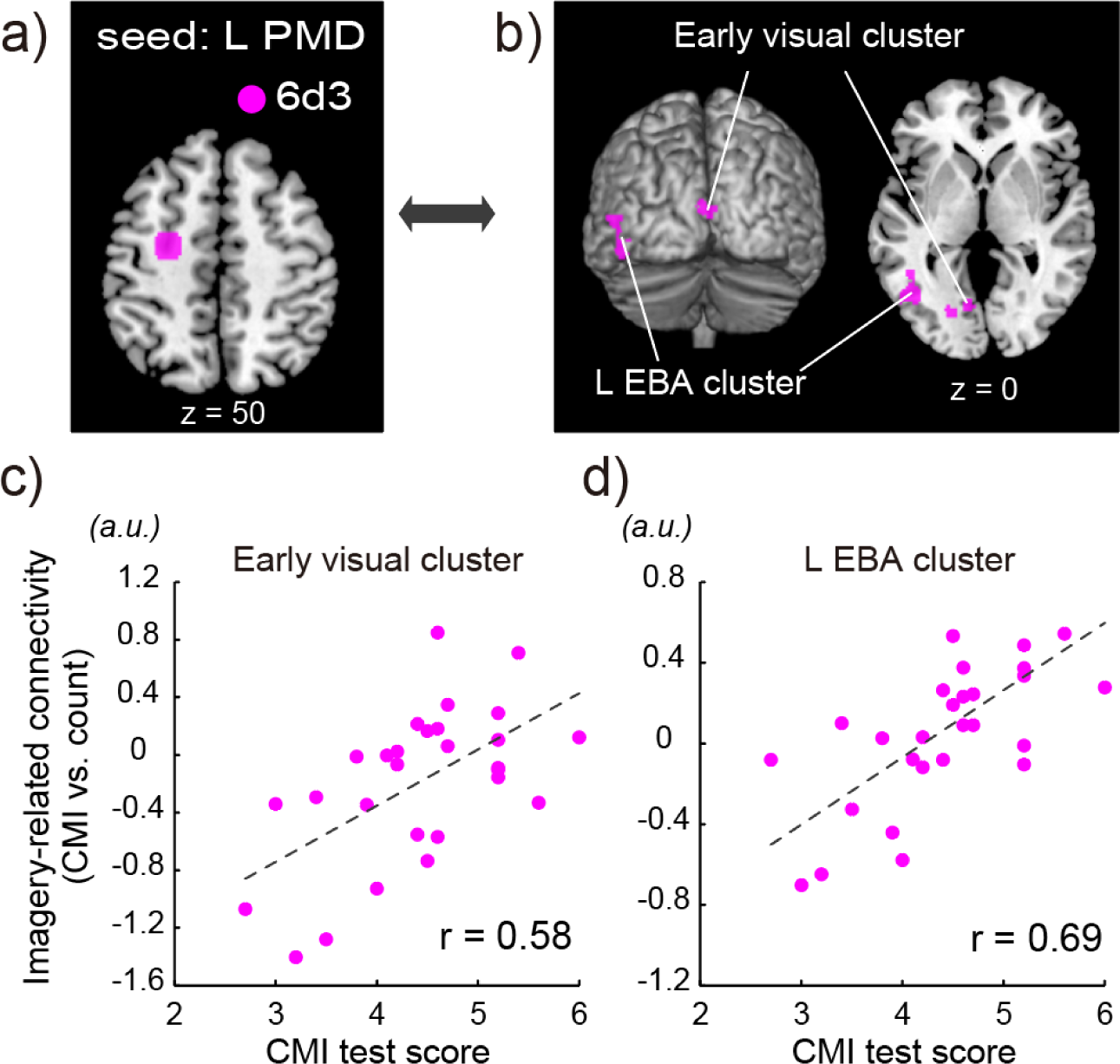
Results of the functional connectivity analysis from the left PMD.**a)** A seed region in the left PMD (cytoarchitectonic area 6d3, pink section) superimposed on the horizontal section (z = 50) of the MNI standard brain. **b)** The left EBA and left- early visual clusters of voxels whose imagery-related connectivity with the left PMD positively correlated with the CMI test score. These are superimposed on the MNI standard brain and its horizontal section (z = 0). **c, d)** Interindividual correlation between the imagery-related connectivity (vertical axis) of the early visual (panel c) or EBA (panel d) cluster and the CMI test score (horizontal axis). Each dot represents individual data. Abbreviations: EBA, extrastriate body area; MNI, Montreal Neurological Institute.

When the right area 5L was a seed region (Figure 5a), we found five significant clusters in the left PMD and primary sensorimotor cortices (PMD–SM1 arm section), left secondary somatosensory cortex (parietal opercular region), and higher-order visual cortices (EBA and fusiform body area [FBA]; Figure 5b). For each cluster, we plotted the relationship between the CMI test score and the imagery-related connectivity for all participants and found a correlation coefficient of r = 0.56 for the left PMD–SM1 cluster, r = 0.63 for the left parietal opercular cluster, r = 0.61 for the left EBA cluster, r = 0.62 for the right EBA cluster, and r = 0.78 for the right FBA cluster (Figure 5c–5g). When the right area 7A was a seed region (Figure 5a), we found two significant clusters in the bilateral EBA, which were closely located to those identified in the above (right area 5L) analysis (Figure 5b). The correlation coefficient between the CMI test score and the imagery-related connectivity across all participants was r = 0.76 for the left EBA cluster and r = 0.51 for the right EBA cluster (Figure 5h, 5i).

**Figure 5.**
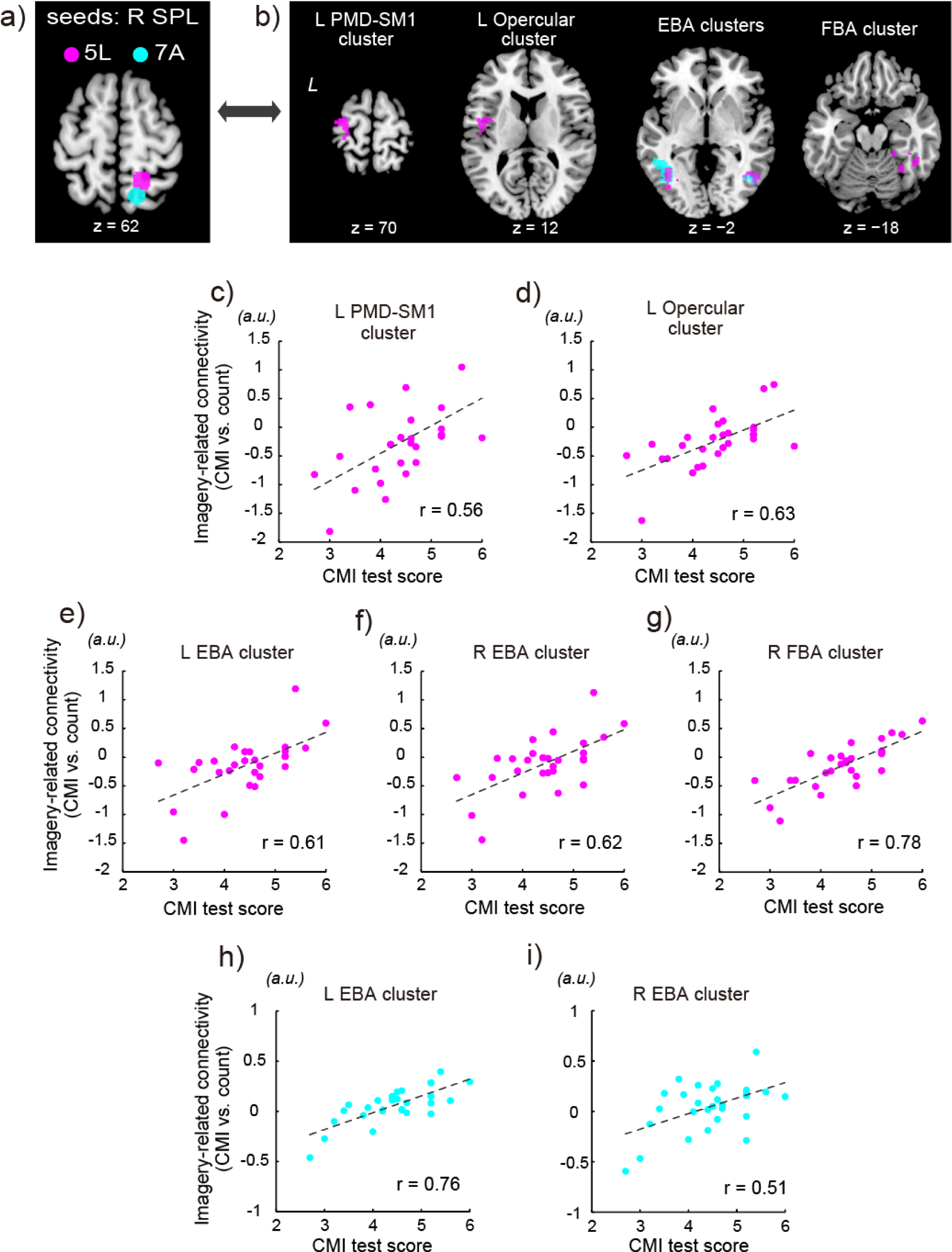
Results of the functional connectivity analysis from the right SPL.**a)** Seed regions in the right area 5L (pink section) and right area 7A (light blue section) are superimposed on the horizontal section (z = 62) of the MNI standard brain. b) The left PMD-SM1, left parietal opercular, bilateral EBA, and right FBA clusters (pink sections), whose imagery-related connectivity with the right area 5L was correlated with the CMI test score, are superimposed on the horizontal section (z = 70, 12, −2, and −18) of the MNI standard brain, respectively. The bilateral EBA clusters (light blue sections), whose imagery-related connectivity with the right area 7A correlated with the CMI test score, are also superimposed on the horizontal section (z = −2). **c–g)** Imagery-related connectivity from the right area 5L. Interindividual correlation between the imagery-related connectivity (vertical axis) of the PMD-SM1 (panel c), parietal opercular (panel d), left EBA (panel e), right EBA (panel f), or right FBA (panel g) cluster and the CMI test score (horizontal axis). **h, i)** Imagery-related connectivity from the right area 7A. Interindividual correlation between the imagery-related connectivity (vertical axis) of the left EBA (panel h) or the right EBA (panel i) cluster and the CMI test score (horizontal axis). Each dot represents individual data. Abbreviations: EBA, extrastriate body area; FBA, fusiform body area; MNI, Montreal Neurological Institute; PMD, dorsal premotor cortex; SM1, primary somatosensorimotor cortices; SPL, superior parietal lobule.

When the left area 5L was a seed region, we found a significant cluster in higher-order visual cortices (not EBA and FBA, not shown in figure), and when the left area 7A was a seed region, no significant clusters were found. Thus, during the CMI task, mainly the left PMD and/or the right SPL increased their functional coupling with the early (primary) and higher-order visual, somatosensory, and motor cortices in higher scorers on the CMI test.

Importantly, none of the regions in which imagery-related connectivity positively correlated with the CMI test score showed a significant increase in the imagery-related activity during the CMI task (Figure 3). Thus, only functional coupling increased in these regions, not the activity itself.

## Discussion

### Objective evaluation of motor imagery by the CMI test

The subjective rating of confidence was not directly related to individual controllability of motor imagery objectively evaluated by the CMI test. This means that those who reported feeling confident during the CMI task do not necessarily exhibit higher CMI test scores. In the present study, the Japanese version of the MIQ-R^15^ (JMIQ- R), in addition to the CMI test, were also administered in 24 out of 89 participants. In the JMIQ-R test, participants gave subjective rating on a scale of 1–7 (kinesthetic score) of how easy to imagine a certain action as if they were actually performing it after performing the action. The mean score of the 24 participants on the CMI test was 4.8 ± 0.7 (range 3.3–5.9), the mean kinesthetic score on the JMIQ-R test was 6.3 ± 0.7 (range 4.8–7.0), and these were not significantly correlated (r = −0.15, N = 24, p = 0.49). These results indicate that subjective rating of motor imagery, which has been often used in previous studies, may not be directly related to the individual’s capability to manipulate motor imagery that can be objectively assessed by the CMI test.

### White matter expansion in the SLF in individuals with higher scores on CMI test

The white matter regions where the SLF is likely running expanded in individuals with higher CMI test scores (Figure 2a–2c), suggesting that individuals with expanded SLF are good at motor imagery (=have higher controllability of motor imagery). To date, neuroimaging studies on motor imagery have mainly focused on its functional neural correlates^6^. To our knowledge, the present study is the first to elucidate the brain structures characterized in individuals who have higher controllability of motor imagery. We originally expected white matter expansion where SLF I or II is likely running. However, we also found the expansion where SLF III is likely running. SLF III most likely connects the inferior frontoparietal cortices^27–29^, and the meta-analysis of motor imagery study^6^ showed that involvement of the inferior frontoparietal cortices in motor imagery. Indeed, we also found imagery-related activity in the right IPL. Thus, white matter expansion where SLF III is likely running may be a reasonable finding.

Although the exact physiological changes underlying white matter expansion are not fully understood, white matter changes are thought to be related to changes in the number of axons, axon diameter, packing density of fibers, axon branching, axon trajectories, and myelination^38^. Thus, if we may assume that the present white matter expansion reflects SLF expansion, the white matter expansion in higher scorers suggests fast and rich information processing in their frontoparietal network connected by the tract. Studies have proposed that white matter tracts develop in an activity-dependent manner^39–41^. As described in the Introduction, the superior frontoparietal regions connected by SLF I or II play a preferential role in spatial/motor processing, and the inferior frontoparietal regions connected by SLF III (particularly the right side) play a dominant role in self-body-related information processing^42,43^. In this study, persons with sports training experiences showed higher CMI test scores than non-experienced ones, which replicates the previous report^21^. Sports experiences that most likely require spatial/motor and corporeal processing could be one of the factors to SLF development in general.

### Imagery-related activity during the CMI test

As we hypothesized, imagery-related activity during the CMI task was identified in bilateral PMD and SPL, which are likely connected by SLF I or II (Figure 3). This suggests the importance of this network for motor imagery, which is consistent with previous reports (Introduction). Bilateral PMD–SPL network is the core network for spatial processing, e.g., spatial imagery (mental rotation) and spatial working memory^44^. During the CMI task, one has to generate, manipulate, and hold one’s imaginary postures through a trial; thus, mental simulation during the CMI task requires the above spatial cognitive functions. One may argue that the present PMD–SPL activity merely reflects the working memory component. However, the PMD–SPL network has been reported to become active during motor imagery tasks in which working memory component is not required^31,32,35,36^. Taken together, we may assume that the PMD–SPL network can be considered the core network for mental simulation (manipulation) based on the spatial cognitive functions.

A previous functional MRI (fMRI) decoding study^45^ demonstrated that two types of hand/finger movements can be classified from the preparatory activity in the PMD. If we may consider motor imagery an extended version of motor preparation process^1^, the present PMD activity may conceivably contain information of the movement contents to be imagined, which is essential for mental simulation of movements. On the contrary, a previous study^45^ also showed that the movements can be classified from the SPL activity only during the execution period when motor commands, efference copy, and sensory feedback are processed in the brain, suggesting that the SPL plays distinct roles (presumably state estimation^46^) from the PMD during motor imagery, although they are forming a functional network (see also below).

Imagery-related activity was also found in the left LOC, left precuneus, and right IPL. Of these, the left LOC appears to correspond to Brodman area 39, which is a constituent of the network connected by SLF I or II^27^. In this study, we confirmed that the left LOC region is likely a constituent of the network connected by SLF I or II (Figure 3a, 3b). In the CMI task, participants were required to imagine in first person with their eyes closed, as if they were actually performing the movement as instructed. Such first-person motor imagery is often treated as kinesthetic motor imagery; however, such imagery also contains a visual component^22^. The left LOC and precuneus are brain structures activated when participants mentally visualize hand movements from the first-person perspective with no actual movements^47^. Thus, these regions could be associated with visual motor imagery from the first-person perspective.

The imagery-related activity in the right IPL is also worth discussing (Figure 3c). Activity in this region was also reported in a meta-analysis^6^. This region corresponds well to the region involved in proprioceptive (kinesthetic) information processing from all limbs, regardless of the left or right limb^48^, and to the region involved in proprioceptive (position) matching between the left and right lower limbs^49^. This region is a constituent of the inferior frontoparietal network connected by SLF III^28^ as we confirmed in the present study (Figure 3c), and its activity is associated with kinesthetic awareness^28^. Indeed, electrical stimulation to this region elicits sensation of illusory limb movement^50^. Furthermore, similar region was found to become active when participants view their actions from a first-person perspective^51^. Thus, speculatively, the right IPL activity could be related to participant’s first-person kinesthetic and visual motor imagery, although the brain does not receive real kinesthetic and visual inputs during the CMI task. Another possible interpretation would be that the right IPL activity was associated with the conscious intention to generate motor imagery as electrical stimulation to this region was also reported to elicit sensation of “desire to move limbs”^50,52^.

Although previous studies have reported the activation of the supplementary motor area (SMA), cerebellum, basal ganglia, and other areas associated with motor imagery^6^, no imagery-related activities were identified in these areas in the present study. The lack of imagery-related activities in these areas could be due to the current experimental design. In the present study, we used a count task as a control task. A previous study using a task similar to the present count task reported SMA and cerebellar activities during the task^53^. In the present study, we also confirmed activity increases in these regions during the count task (not shown in the figure). Although these regions are inherently involved in motor imagery, any further significant increase in activity during the CMI task was not detected in the present between-condition comparison design.

### Imagery-related functional connectivity

The left PMD (area 6d3) increased the imagery-related connectivity with the left- early and higher-order visual cortices in higher scorers on the CMI test (Figure 4). Similarly, in higher scorers, the right area 5L increased the connectivity with the left PMD–SM1, left secondary somatosensory cortex, and bilateral higher-order visual cortices, and the right area 7A increased the connectivity with the bilateral higher-order visual cortices (Figure 5).

All higher-order visual cortices described above correspond to the EBA and FBA, which preferentially respond to visual images of body parts or the entire body^54–57^. In addition, regarding the brain regions that showed enhanced functional coupling with the right area 5L, the peaks of the activity in the left secondary somatosensory cortex are located in the rostral parietal opercular region (e.g., cytoarchitectonic area OP4). This region has closer anatomical and functional connections with the premotor cortices and M1^58^ and is proposed to be involved in sensorimotor awareness^59,60^. Furthermore, the peaks of the activity in the left PMD–SM1 are located in the cytoarchitectonic areas 6d1 and 4a, which are involved in kinesthetic processing and motor output^42,61^.

By contrast, no regions showed increased functional coupling with the right PMD. Similarly, the left area 5L increased the connectivity merely with the left higher-order visual cortices (not EBA and FBA), and no regions showed increased functional coupling with the left area 7A. Thus, in the brains of higher scorers on the CMI test (=persons with higher controllability of motor imagery), mainly the left PMD and/or the right SPL increased their functional coupling with the above-mentioned visual body, somatosensory, and motor/kinesthetic areas during motor imagery. These lines of evidence suggest the importance of the left PMD and right SPL in motor imagery among bilateral PMD and SPL, which appears to be compatible with the previous report that motor imagery is compromised in patients with left-frontal and right-parietal damage^23^.

During the CMI test, participants with eyes closed did not actually performed the movements. Thus, the brain did not produce any actual movements nor receive any real visual and somatic inputs normally associated with actual movements. Despite this situation, the left PMD and/or right SPL of higher scorers increased their functional coupling with the visual body, somatosensory, and motor/kinesthetic areas, with no significant increase in imagery-related activity in these areas (Figure 3). Thus, higher controllability of motor imagery requires a top–down access from the left PMD and/or right SPL to these sensory and motor/kinesthetic areas. Although further proof is needed, this access may create more vivid vicarious sensory experiences during motor imagery. This view appears to be compatible with the view of sensory emulation process during motor imagery. The brain emulates various sensory experiences (feedback) that would be expected (predicted) if the imagined movements were actually performed by referring memories of previous enactments^2,5,34,47,62–64.^

The imagery-related connectivity between the PMD–SPL network and the visual body, somatosensory, and motor/kinesthetic areas correlated with the CMI test score (Figures 4 and 5), whereas the imagery-related activity in the PMD–SPL network did not correlate with the score. Thus, the controllability of motor imagery seems appears to be more related to an increase in the functional coupling of activity between the PMD–SPL network and the visual body, somatosensory, and motor/kinesthetic areas, rather than an increase in the activity of the core mental simulation network of PMD and SPL per se. This suggests the importance of sensory emulation for higher controllability of motor imagery and the hub role of the PMD–SPL network in this sensory emulation during motor imagery.

Within the right SPL, the right area 5L increased the functional coupling with the primary and secondary somatosensory and motor areas and with the visual body areas, whereas the right area 7A increased the connectivity only with the visual body areas (Figure 5). The area 7A did not increase the connectivity with somatosensory and motor areas even when a lenient threshold was used (height threshold p < 0.01 uncorrected). Thus, area 5L appears to play a distinct role from area 7A. The former can play a role as a higher-order somatosensory association area, whereas the latter is mainly a higher-order visual association area, which is generally compatible with the view proposed by previous studies^65,66^. We may further raise the possibility that functional roles may differ between the left PMD and the right SPL, i.e., the former may be preferentially involved in visual emulation^47^, whereas the latter was not only visual bodily but also somatosensory and kinesthetic emulation.

Finally, the functional coupling between the right area 5L and the left PMD–M1 deserves discussion, particularly in terms of the motor output (Figure 5b, 5c). Motor imagery of hand movements usually increases the motor–cortical excitability of the hand muscles^67,68^ and generates event-related desynchronization in the SM1^69,70^. In addition, mental practice with motor imagery may expand M1 hand representation^71^, which is a neural basis of early motor skill learning^72,73^. Despite these findings, robust M1 activity is not reported in most neuroimaging studies for motor imagery^6^, except focal activation in area 4a^3,74^. In the present study, we found no significant imagery-related activity in M1 (Figure 3a). However, if we consider that the more realistically one can imagine a movement, the higher the excitability of the M1 for an agonistic muscle of the movement^19^, we may speculate that increased functional coupling between the right area 5L and the left PMD–M1 observed in higher scorers may affect their motor–cortical excitability during motor imagery, which should be proven in future studies.

In conclusion, this study introduced the CMI test that can objectively evaluate an individual’s ability to manipulate one’s imaginary postures and has elucidated structural and functional features that characterize the brains of individuals with higher controllability of motor imagery. Structurally, such individuals have expanded frontoparietal white matter that enables fast and rich neural processing of spatial/motor and corporeal information. Functionally, in their brains, the core network of mental simulation (superior frontoparietal network of PMD–SPL), particularly the left PMD and/or the right SPL, likely has top–down access to the visual body, somatosensory, and motor/kinesthetic areas and enhances functional coupling with these for sensory emulation (prediction). This study advanced the understanding of individual difference in motor imagery ability.

## Methods

### Participants

A total of 89 healthy right-handed young adults (53 men, 36 women: mean age, 22.1 ± 1.7; range, 20–29 years) participated in this study. None of them had a history of self-reported neurological, psychiatric, or motor disorders. Their handedness was confirmed using the Edinburgh handedness inventory^75^. All participants went through the CMI test for the first time. They first performed the CMI test outside the MRI scanner room, and their T1-weighted structural images were acquired. Of the 89 participants, 33 also participated in the fMRI experiment. However, the data obtained from five participants (see below) were excluded, and those obtained from 28 participants (15 men, mean age, 21.6 ± 1.4 years; range, 20–26 years) were analyzed. The study protocol was approved by the Ethics Committee of the National Institute of Information and Communications Technology and the MRI Safety Committee of the Center for Information and Neural Networks (CiNet; no. 2003260010). Details of the experiment were explained to each participant before the experiment, and they provided written informed consent. This study was conducted in accordance with the principles and guidelines of the Declaration of Helsinki (1975).

### CMI test

We employed the CMI test originally introduced by Nishida et al. (1986)^20^. The reliability and validity of this test were evaluated in this study. The reliability was also checked, as we slightly modified the original test in the present study (Cronbach’s alpha coefficient was 0.74). All participants performed 10 trials (sets of questions). One question consisted of six consecutive verbal instructions including the basic posture. The order of the 10 trials (Nos. 1–10) was identical across all participants. After careful explanation and instruction of the CMI test, we started the CMI test.

Each trial consisted of an imagery phase and an answer phase. Figure 1A demonstrates an example of a trial. During the imagery phase, six consecutive verbal instructions about body movements (postures) were provided. Among the instructions, the first instruction was about the basic posture, which was common across all trials. The following five instructions were related to the movements of the body parts (left or right arm, left or right leg, upper body, or head/neck), which was different across trials. An interval between each instruction was approximately 2 s, and the participants listened to the instructions recorded in a computer.

During the imagery phase, starting from the basic posture, the participants had to manipulate their imaginary postures in response to each of five instructions, as if they were actually performing the instructed movements from a first-person perspective, while they closed their eyes and were seated. Thus, during the imagery phase, they had to keep updating their imaginary postures by adding a new posture of one body part for each instruction. No actual body movements were executed, and they were prohibited from remembering the postural changes by any means other than imagery, such as words or other memory techniques.

After the final instruction, a computer-generated “stop” signal was given, and the participants were requested to stand with their eyes open and actually perform the final posture they had in their minds (answer phase)^21^. On the floor, we put guidelines indicating directions in 45° increments and markers indicating distances from the center (50 cm and 1 m). By referring to these indications, they were asked to accurately demonstrate the directions and locations (distances) of body parts in the final posture. In the answer phase, they were allowed to perform only the parts they could answer when they could not answer the entire posture.

The final posture was recorded by a digital video camera and judged offline for correctness. Information of the correct answer was not given to the participants during the CMI test outside of the scanner room. In the offline analysis, a score was given when the final posture (direction, location, and distance) of each body part (left or right arm, left or right leg, upper body, or head/neck) was correct. A person who did not know the purpose of this study consistently evaluated the correctness of the final posture using the same criteria. In each trial, the number of body parts that were answered correctly was counted (maximum score = 6), and the average number of correct answers across the 10 trials was calculated for each participant and used as the CMI test score.

### MRI experiments Structural MRI

After the CMI test outside the scanner room, the participants entered the MRI scanner room and were placed in an MRI scanner. The head was secured with a sponge cushion and adhesive tape, and the participants wore earplugs and MRI-compatible headphones. The arms were set in a natural semi-rotated position, stretched along the body and relaxed. A T1-weighted image was acquired, with a magnetization-prepared rapid gradient echo (MP-RAGE) sequence using a 3.0-Tesla MRI scanner (Trio Tim; SIEMENS, Germany) and a 32-channel array coil for each participant. The imaging parameters were as follows: repetition time [TR], 1,900 ms; echo time [TE], 2.48 ms; flip angle, 9°; field of view, 256 × 256 mm^2^; matrix size, 256 × 256 pixels; slice thickness, 1.0 mm; voxel size = 1 × 1 × 1 mm^3^; contiguous transvers slices, 208.

### fMRI

Of the 89 participants, 33 performed the following fMRI experiment. Functional images were acquired using T2*-weighted gradient echo-planar imaging (EPI), with the same scanner and coil as in the structural images. Each volume consisted of 48 slices (slice thickness, 3 mm) acquired to cover the entire brain using multiband imaging (multiband factor, 3; Moeller et al. 2010). The imaging parameters were as follows: TR, 1000 ms; TE, 30 ms; flip angle, 60°; field of view, 192 × 192 mm^2^; matrix size, 64 × 64 pixels; voxel size, 3 × 3 × 3 mm^3^. In total, 212 volumes were collected per run for both CMI and count tasks.

### Task procedure

Brain activity was scanned while the 33 participants performed a CMI task and a control (count) task. Each participant completed two runs (212 s for each) for each task. The time course of each run is shown in Figure 1b. Each run comprised five task epochs, each lasting 28 s. Each epoch was separated by 12-s baseline (rest) periods. Each run also included a 12-s baseline period before the start of the first epoch.

In the CMI task (Figure 1b), the participants performed the same imagery task as the CMI test performed outside the scanner room. Each epoch corresponded to the imagery phase of each trial in the CMI test. The first instruction about the basic posture in each epoch (during the imagery phase in each trial) was eliminated, as this was common across all trials. Even though the first instruction was eliminated, they were explicitly instructed to imagine body-part movements as per instructions from the basic posture in each epoch. In the CMI task, the order of trial was changed from the CMI test to eliminate the possible effect of participants remembering the order of the trials performed in the CMI test. The trials (Nos. 2, 10, 8, 9, and 1) were assigned in a run in this order (run A), whereas the remaining trials (Nos. 5, 7, 6, 3, and 4) was assigned in another run (run B).

At the end of each epoch, after the final instruction was completed, a computer-generated “stop” signal was given. In the answer phase following each epoch, since the final posture cannot be performed in the scanner, we asked the participants to rate how confident they were about the accuracy of their imaginary final posture (subjective rating of confidence). They were asked to press a corresponding button with their right fingers on a four-point scale (1, not at all; 2, not very well; 3, fairly well; 4, very well) within approximately 3 s after the “stop” signal. All instructions were given audibly through MRI-compatible headphones, and the responses were recorded using an MR-compatible four-button device (Current Design Inc., Philadelphia, PA).

In the count task, the participants listened to the same instructions as in the CMI task in each epoch. However, they were asked not to imagine body movements. Instead, they had to count the total number of the term “right” (e.g., right arm, right leg, and rightward), which appeared during five instructions in each epoch. At the end of each epoch, after the final instruction was completed, a computer-generated “stop” signal was given. Within approximately 3 s after the “stop” signal, they had to answer the total number by pressing a corresponding button with their right fingers. Since the total number ranged from 1 to 4 across all trials, they had to press a button either from 1 to 4 on the same four-button device. Thus, in this control task, the participants must pay attention to and process the same verbal instructions; however, its associated mental process was different from the mental imagery required during the CMI task. Thus, by comparing brain activity during the CMI task with that during the counting task, one may identify brain activity associated with mental imagery (manipulation) processes (imagery-related activity).

To counterbalance the task order, we prepared a group that performed the CMI task first and a group that performed the count task first. In addition, to counterbalance the order of runs, we assigned a group that performed run A first and a group that performed run B first for both the CMI and count tasks. Eventually, four groups were arranged. Participants who made three or more errors on the correct “right” count in the count task were considered to lack concentration on the instructions and were excluded from the analysis. Participants were added until the number of participants with less than three errors reached seven in each group. Indeed, all 28 participants made no more than one error.

### MRI data analysis

#### Structural MRI analysis: voxel-based morphometry (VBM)

Using a VBM analysis, we searched for brain regions (both white and gray matter) whose tissue volume correlated with the CMI test scores to examine the morphological features of the brains in individuals with higher ability to manipulate motor imagery.

We first visually inspected the T1-weighted structural images of all participants and confirmed the absence of observable structural abnormalities and motion artifacts. Data were then analyzed using Statistical Parametric Mapping (SPM 12, Wellcome Center for Human Neuroimaging, London, UK) running on MATLAB R2018b (Math Works, Sherborn, MA, USA). The following analysis procedures were conducted as recommended by Ashburner (2010)^76^ and used in our previous study^77^.

First, the T1-weighted structural image of each participant was segmented into gray matter, white matter, cerebrospinal fluid (CSF), and non-brain parts based on a tissue probability map provided by SPM. Through this procedure, segmented gray and white matter images were generated. These images approximated to the tissue probability map by rigid body transformation, and transformed gray matter and white matter images were generated.

Then, diffeomorphic anatomical registration through exponentiated lie algebra (DARTEL) processing was performed to generate a DARTEL template, which was later used to accurately align the segmented images across participants. In this process, the average image of the transformed gray and white matter images across all participants was defined as a template that includes the gray and white matter. A deformation field was calculated to deform the template into the transformed images of each participant, and the inverse of the deformation field was applied back to the individual images. To eventually generate a sharply defined DARTEL template, this series of operations were performed multiple times.

Thereafter, an affine transformation was applied to the DARTEL template to align it with the tissue probability map in the Montreal Neurological Institute (MNI) standard space. Then, the segmented gray and white matter images of each participant were warped nonlinearly to the DARTEL template represented in the MNI space (spatial normalization). The warped image was modulated by Jacobian determinants of the deformation field to preserve the relative gray and white matter volume, even after spatial normalization.

Finally, the modulated gray and white matter images of each participant was smoothed with a 6-mm full-width-at-half-maximum (FWHM) Gaussian kernel and resampled to a resolution of 1.5 × 1.5 × 1.5 mm voxel size.

In the group-level analysis, to identify brain regions in which the tissue volume correlated with the CMI test score, multiple regression analysis was performed for the white and gray matter. Individual CMI test scores were included as independent variables, as well as sex, age, and intracerebral volume (sum of the gray matter, white matter, and CSF volumes) as nuisance covariates (i.e., effects of no interest) because these factors may have a significant effect on the results of a VBM analysis^78^. To exclusively select truly positive white or gray matter voxels by eliminating possible noise outside the brain and increase statistical power by reducing search volumes, we generated a mask based on our present data using the SPM Masking Toolbox^79^. Thus, voxels outside this mask were excluded from the analysis. We identified active voxels using a height threshold of p < 0.005 and an extent threshold of p < 0.05, corrected for multiple comparisons with the family-wise error rate across the entire brain. This threshold was consistently used in the subsequent functional analysis.

In the white matter region, we found a significant cluster of voxels that showed positive correlation with the CMI test score in each of the left and right hemispheres. To verify that the bilateral white matter clusters overlap with the bilateral SLF (I, II, or III), we used a simple atlas-based overlay approach^28^. We visualized the overlap between the white matter clusters and the tract probability maps of SLF I, II, or III using the MRIcron software (Figure 2a–2c) and calculated the volume of the overlapping regions. For this, we used tract probability maps depicting SLF I, II, or III obtained from publicly available databases (https://storage.googleapis.com/bcblabweb/open_data.html), with the threshold of 50% probability. Each tract probability map of SLF I, II, or III describes the stream of each fiber tract. However, the maps of SLF I, II, or III had overlapped sections among them in each hemisphere. Then, we calculated the overlapping volume between the white matter clusters and each of the pure SLF I, II, or III and the overlapping volume between the white matter clusters and overlapped sections among SLF I, II, and III, separately (Figure 2d). This was a purely descriptive approach where we did not perform any statistical evaluation.

To confirm positive correlation between the parameter estimates of each cluster and CMI test scores, we extracted the parameter estimates from all voxels constituting each cluster for each participant and calculated their means for each cluster. Then, we plotted the parameter estimates against the CMI test scores across all participants for each cluster, after removing the effects of age, sex, and total brain volume. To avoid circular statistical evaluation, we simply displayed the correlation between them and its coefficient (Figure 2e, 2f).

#### fMRI analysis Preprocessing

To eliminate the effect of unsteady magnetization during the tasks, the first four EPI images in each run were discarded. The functional imaging data were analyzed using SPM 12 running on MATLAB R2018b.

First, we corrected for head motion caused by body movements and heartbeats during the experiment (realignment). All EPI images were aligned to the first EPI image of the first session using six degrees of freedom (translation and rotation about the X-, Y-, and X-axes) for rigid displacement. Through this realignment procedure, time series data of the head position during the scanning were obtained. To evaluate head motions, we calculated the absolute value of displacement in each frame from its previous frame (framewise displacement [FD])^80^. The number of frames in which FD exceeded 0.9 mm was 0 in most participants (26/28). Even in participants who had such frames, the percentage was <1% of all 832 frames. Therefore, we excluded no data from the analysis. Then, the T1-weighted structural image of each participant was coregistered to the mean image of all realigned EPI images using affine transformation. Then, the structural and realigned EPI images were spatially normalized to the standard MNI space^81^. Normalization parameters for aligning the structural image with the MNI template brain were calculated using the SPM12 normalization algorithm. The same parameters were used to transform the realigned EPI images. Finally, the normalized images were spatially smoothed using a 6-mm FWHM Gaussian kernel.

#### Single subject analysis

We analyzed the preprocessed imaging data using a general linear model (GLM)^82,83^. A design matrix is prepared for each participant. The design matrix included a boxcar function for a task epoch in a run, which was convolved with the hemodynamic response function (HRF). To account for the effect of button press on brain activity, an impulse function was assigned to the timing of each button press, which was convolved with the HRF. Finally, to correct the residual motion-related variance after realignment, six realignment parameters were included in the design matrix as regressors of no interest. For each participant, a contrast image showing brain activity during the CMI task (CMI task > baseline) was created. To specify imagery-related activity, a contrast image showing greater activity during the CMI task than during the count task (CMI task > count task) was created.

### Group analysis

Contrast images from all participants were entered into a second-level random-effects group analysis^84^. A one-sample t-test was performed for the contrast image showing imagery-related activity (CMI task > count task). In this analysis, a cluster image that showed an increase in activity (height threshold p < 0.05 uncorrected) during the CMI task (CMI task > baseline) was used as an inclusive mask. By using this mask image at the liberal threshold, we ensured that observed imagery-related activation is true activity rather than pseudo-activation caused merely by deactivation in the count task. To identify anatomical regions of activation, we referred to the cytoarchitectonic probability maps implemented in the JuBrain Anatomy toolbox v3.0^85,86^.

To verify that the frontoparietal activations are located in the cortical regions with which SLF I, II, or III is likely connecting, we used the same overlay approach used in the VBM analysis (Figure 3a–3c). Each tract probability map of SLF I, II, or III not only describes the stream of each fiber tract in the white matter but also its connecting cortical regions. As we did in the VBM analysis, we simply reported overlapping volume between the frontoparietal activations and cortical regions with which SLF I, II, or III is likely connecting (see above). This was again a purely descriptive approach where we did not perform any statistical evaluation.

### Functional connectivity analysis

Finally, we examined brain regions where imagery-related functional coupling with a seed region changes in relation to the CMI test score across participants by conducting a generalized psychophysiological interaction analysis (gPPI)^87^. This analysis was performed on preprocessed fMRI data using the CONN toolbox version 21a^88^. Physiological noise originating from the white matter and CSF were removed using the component-based noise correction method (CompCor) in the toolbox^89^. A temporal band- pass filter of 0.009–0.08 Hz was applied.

We prepared eight seed regions based on the peaks of the imagery-related activity in the bilateral PMD–SPL networks (Figure 3). Each seed region was defined as a spherical region of 8-mm radius centered at the peak coordinates. In the left PMD, we prepared two seed regions in areas 6d3 (peak coordinates: x = −26, y = −8, z = 48) and 6d1 (peak coordinates: −16, −6, 70). Similarly, in the right PMD, we prepared two seed regions in areas 6d3 (peak coordinates: 24, −6, 54) and 6d1 (peak coordinates: 20, −8, 64). As for the SPL in each hemisphere, we prepared a seed region in each of areas 5 and 7 because the former can play a role as higher-order somatosensory association area, whereas the latter as higher-order visual association area^65,66^. In the left SPL, we prepared two seed regions in areas 5L (peak coordinates: −22, −48, 68) and 7A (peak coordinates:−18, −54, 60). In the right SPL, we prepared two seed regions in areas 5L (peak coordinates: 18, −50, 60) and area 7A (peak coordinates: 14, −62, 58). We confirmed that all peaks were in the cortical regions with which SLF I is likely connecting.

In the gPPI analysis, we used each of the eight seed regions. In each participant, the time course of the average fMRI signal across the voxels in each seed region was deconvolved using the canonical HRF (physiological variable). Then, we performed a general linear model analysis using the design matrix and included the following regressors: physiological variable, boxcar function for the task epoch (psychological variable), and multiplication of the physiological variable and the psychological variable (PPI). These variables were convolved with a canonical HRF. Six realignment parameters were also included in the design matrix as regressors of no interest.

In each task, we first generated an image of voxels showing to what extent their activities changed with the PPI regressor of each seed region in each participant. Then, we generated a contrast image (CMI task > count task) that shows imagery-related connectivity change for each participant. We used this individual image in the second-level group analysis, in which we examined brain regions where imagery-related connectivity changes in relation to the CMI test score across participants. In this analysis, individual CMI scores were included in the design matrix as independent variables. The task and run orders were also included as nuisance covariates, to exclude the possibility that these factors affect the results since these orders were counterbalanced across participants.

We examined significant clusters throughout the brain (Figure 4b and 5b). When a significant cluster was identified, we confirmed positive correlation between the parameter estimates of each cluster and the CMI test scores. We extracted the parameter estimate of the imagery-related connectivity from all voxels constituting each cluster for each participant, and calculated the mean of them for each cluster. We then plotted the parameter estimates against the CMI test scores across all participants for each cluster, after removing the effects of age, gender, and total brain volume. To avoid circular statistical evaluation, we simply displayed the correlation between them and its coefficient (Figure 4c–d, 5c–i).

### Data availability

The data that support the findings of this study are available from the corresponding author upon reasonable request.

## Acknowledgments

This study was supported by JSPS KAKENHI Grant Nos. JP19H05723, JP23H03706, and JP23K17453 for EN and JSPS KAKENHI Grant No. JP20H04492 for TM. The funding sources were not involved in the study design; collection, analysis, interpretation of data; writing of the report; or the decision to submit the article for publication. The authors are grateful to Dr Tsuyoshi Ikegami, Dr Jihoon Park, and Dr Hideki Nakano for their valuable comments on this work. They also thank CiNet MRI staff for their support, Ms Keiko Ueyama for the illustration, and Mr Susumu Minamiyama for helping with the data analysis.

## Author contributions

All authors contributed to the conception and study design, data collection or acquisition (TM and EN), statistical analysis, interpretation of results, drafting the manuscript work or revising it critically for important intellectual content, and approval of final version to be published and agreement to be accountable for the integrity and accuracy of all aspects of the work.

## References

1. Jeannerod, M. The representing brain: Neural correlates of motor intention and imagery. Behav. Brain Sci. 17, 187–202; 10.1017/S0140525X00034026 (1994).

2. Naito, E. et al. Internally simulated movement sensations during motor imagery activate cortical motor areas and the cerebellum. J. Neurosci. 22, 3683–3691; 10.1523/JNEUROSCI.22-09-03683.2002 (2002).

3. Ehrsson, H. H., Geyer, S. & Naito, E. Imagery of voluntary movement of fingers, toes, and tongue activates corresponding body-part-specific motor representations. J. Neurophysiol. 90, 3304–3316; 10.1152/jn.01113.2002 (2003).

4. Hanakawa, T., Dimyan, M. A. & Hallett, M. Motor planning, imagery, and execution in the distributed motor network: A time-course study with functional MRI. Cereb. Cortex 18, 2775–2788; 10.1093/cercor/bhn036 (2008).

5. Hanakawa, T. Organizing motor imageries. Neurosci. Res. 104, 56–63; 10.1016/j.neures.2015.11.003 (2016).

6. Hétu, S. et al. The neural network of motor imagery: An ALE meta-analysis. Neurosci. Biobehav. Rev. 37, 930–949; 10.1016/j.neubiorev.2013.03.017 (2013).

7. Mizuguchi, N., Nakata, H., Uchida, Y. & Kanosue, K. Motor imagery and sport performance. J. Phys. Fit. Sports Med. 1, 103–111; 10.7600/jpfsm.1.103 (2012).

8. Martin, K. A., Moritz, S. E. & Hall, C. R. Imagery use in sport: A literature review and applied model. Sport Psychol. 13, 245–268; 10.1123/tsp.13.3.245 (1999).

9. Munzert, J. & Zentgraf, K. Motor imagery and its implications for understanding the motor system. Prog. Brain Res. 174, 219–229; 10.1016/S0079-6123(09)01318-1 (2009).

10. Stockley, R. C., Jarvis, K., Boland, P. & Clegg, A. J. Systematic review and meta-analysis of the effectiveness of mental practice for the upper limb after stroke: Imagined or real benefit? Arch. Phys. Med. Rehabil. 102, 1011–1027; 10.1016/j.apmr.2020.09.391 (2021).

11. Ladda, A. M., Lebon, F. & Lotze, M. Using motor imagery practice for improving motor performance-a review. Brain Cogn. 150, 105705; 10.1016/j.bandc.2021.105705 (2021).

12. Isaac, A., Marks, D. & Russell, D. An instrument for assessing imagery of movement: The vividness of movement imagery questionnaire (VMIQ). J. Ment. Imagery 10, 23–30 (1986).

13. Hall, C. & Pongrac, J. Movement imagery questionnaire, (1983).

14. Ziv, G., Lidor, R., Arnon, M. & Zeev, A. The vividness of movement imagery questionnaire (VMIQ-2)-translation and reliability of a Hebrew version. ISR J Psychiatr*y***54**, 48–52 (2017).

15. Hall, C. R. & Martin, K. A. Measuring movement imagery abilities: A revision of the movement imagery questionnaire. J. Ment. Imagery 21, 143–154 (1997).

16. Gregg, M., Hall, C. & Butler, A. The MIQ-RS: A suitable option for examining movement imagery ability. Evid. Based Complement. Alternat. Med. 7, 249–257; 10.1093/ecam/nem170 (2010).

17. Collet, C., Guillot, A., Lebon, F., MacIntyre, T. & Moran, A. Measuring motor imagery using psychometric, behavioral, and psychophysiological tools. Exerc. Sport Sci. Rev. 39, 85–92; 10.1097/JES.0b013e31820ac5e0 (2011).

18. Guillot, A. et al. Functional neuroanatomical networks associated with expertise in motor imagery. Neuroimage 41, 1471–1483; 10.1016/j.neuroimage.2008.03.042 (2008).

19. Lebon, F., Byblow, W. D., Collet, C., Guillot, A. & Stinear, C. M. The modulation of motor cortex excitability during motor imagery depends on imagery quality. Eur. J. Neurosci. 35, 323–331 (2012).

20. Nishida, T. et al. A new test for controllability of motor imagery : The examination of its validity and reliability. *Jpn*. J. Phys. Educ. 31, 13–22 (1986).

21. Naito, E. Controllability of motor imagery and transformation of visual imagery. Percept. Mot. Skills 78, 479–487; 10.2466/pms.1994.78.2.479 (1994).

22. Amemiya, K. et al. Neurological and behavioral features of locomotor imagery in the blind. Brain Imaging Behav. 15, 656–676; 10.1007/s11682-020-00275-w (2021).

23. Johnson, S. H. Imagining the impossible: Intact motor representations in hemiplegics. NeuroReport 11, 729–732; 10.1097/00001756-200003200-00015 (2000).

24. Sirigu, A. et al. The mental representation of hand movements after parietal cortex damage. Science 273, 1564–1568; 10.1126/science.273.5281.1564 (1996).

25. McInnes, K., Friesen, C. & Boe, S. Specific brain lesions impair explicit motor imagery ability: A systematic review of the evidence. Arch. Phys. Med. Rehabil. 97, 478–489.e1; 10.1016/j.apmr.2015.07.012 (2016).

26. Johnson, S. H., Sprehn, G. & Saykin, A. J. Intact motor imagery in chronic upper limb hemiplegics: Evidence for activity-independent action representations. J. Cogn. Neurosci. 14, 841–852; 10.1162/089892902760191072 (2002).

27. Thiebaut de Schotten, M., Dell’Acqua, F., Valabregue, R. & Catani, M. Monkey to human comparative anatomy of the frontal lobe association tracts. Cortex 48, 82–96; 10.1016/j.cortex.2011.10.001 (2012).

28. Amemiya, K. & Naito, E. Importance of human right inferior frontoparietal network connected by inferior branch of superior longitudinal fasciculus tract in corporeal awareness of kinesthetic illusory movement. Cortex 78, 15–30;10.1016/j.cortex.2016.01.017 (2016).

29. Parlatini, V. et al. Functional segregation and integration within fronto-parietal networks. Neuroimage 146, 367–375; 10.1016/j.neuroimage.2016.08.031 (2017).

30. Stephan, K. M. et al. Functional anatomy of the mental representation of upper extremity movements in healthy subjects. J. Neurophysiol. 73, 373–386; 10.1152/jn.1995.73.1.373 (1995).

31. Gerardin, E. et al. Partially overlapping neural networks for real and imagined hand movements. Cereb. Cortex 10, 1093–1104; 10.1093/cercor/10.11.1093 (2000).

32. Hanakawa, T. et al. Functional properties of brain areas associated with motor execution and imagery. J. Neurophysiol. 89, 989–1002; 10.1152/jn.00132.2002 (2003).

33. Kuhtz-Buschbeck, J. P. et al. Effector-independent representations of simple and complex imagined finger movements: A combined fMRI and TMS study. Eur. J. Neurosci. 18, 3375–3387; 10.1111/j.1460-9568.2003.03066.x (2003).

34. Solodkin, A., Hlustik, P., Chen, E. E. & Small, S. L. Fine modulation in network activation during motor execution and motor imagery. Cereb. Cortex 14, 1246–1255; 10.1093/cercor/bhh086 (2004).

35. Lorey, B. et al. Neural simulation of actions: Effector-versus action-specific motor maps within the human premotor and posterior parietal area? Hum. Brain Mapp. 35, 1212–1225; 10.1002/hbm.22246 (2014).

36. Ogawa, T., Shimobayashi, H., Hirayama, J. I. & Kawanabe, M. Asymmetric directed functional connectivity within the frontoparietal motor network during motor imagery and execution. Neuroimage 247, 118794; 10.1016/j.neuroimage.2021.118794 (2022).

37. Sigl, B., et al. The human dorsal premotor cortex-cytoarchitecture, maps and function in OHBM (2016).

38. Zatorre, R. J., Fields, R. D. & Johansen-Berg, H. Plasticity in gray and white: Neuroimaging changes in brain structure during learning. Nat. Neurosci. 15, 528–536; 10.1038/nn.3045 (2012).

39. Fields, R. D. A new mechanism of nervous system plasticity: Activity-dependent myelination. Nat. Rev. Neurosci. 16, 756–767; 10.1038/nrn4023 (2015).

40. Sampaio-Baptista, C. & Johansen-Berg, H. White matter plasticity in the adult brain. Neuron 96, 1239–1251; 10.1016/j.neuron.2017.11.026 (2017).

41. Morita, T., Takemura, H. & Naito, E. Functional and structural properties of interhemispheric interaction between bilateral precentral hand motor regions in a top wheelchair racing Paralympian. Brain Sci. 13, 715; 10.3390/brainsci13050715 (2023).

42. Naito, E., Morita, T. & Amemiya, K. Body representations in the human brain revealed by kinesthetic illusions and their essential contributions to motor control and corporeal awareness. Neurosci. Res. 104, 16–30; 10.1016/j.neures.2015.10.013 (2016).

43. Morita, T. et al. Self-face recognition shares brain regions active during proprioceptive illusion in the right inferior fronto-parietal superior longitudinal fasciculus III network. Neuroscience 348, 288–301; 10.1016/j.neuroscience.2017.02.031 (2017).

44. Cona, G. & Scarpazza, C. Where is the ‘where’ in the brain? A meta-analysis of neuroimaging studies on spatial cognition. Hum. Brain Mapp. 40, 1867–1886; 10.1002/hbm.24496 (2019).

45. Nambu, I. et al. Decoding sequential finger movements from preparatory activity in higher-order motor regions: A functional magnetic resonance imaging multi-voxel pattern analysis. Eur. J. Neurosci. 42, 2851–2859; 10.1111/ejn.13063 (2015).

46. Takei, T., Lomber, S. G., Cook, D. J. & Scott, S. H. Transient deactivation of dorsal premotor cortex or parietal area 5 impairs feedback control of the limb in macaques. Curr. Biol. 31, 1476–1487.e5; 10.1016/j.cub.2021.01.049 (2021).

47. Guillot, A. et al. Brain activity during visual versus kinesthetic imagery: An fMRI study. Hum. Brain Mapp. 30, 2157–2172; 10.1002/hbm.20658 (2009).

48. Naito, E. et al. Human limb-specific and non-limb-specific brain representations during kinesthetic illusory movements of the upper and lower extremities. Eur. J. Neurosci. 25, 3476–3487; 10.1111/j.1460-9568.2007.05587.x (2007).

49. Iandolo, R. et al. Neural correlates of lower limbs proprioception: An fMRI study of foot position matching. Hum. Brain Mapp. 39, 1929–1944; 10.1002/hbm.23972 (2018).

50. Desmurget, M. et al. Movement intention after parietal cortex stimulation in humans. Science 324, 811–813; 10.1126/science.1169896 (2009).

51. Asakage, S. & Nakano, T. The salience network is activated during self-recognition from both first-person and third-person perspectives. Hum. Brain Mapp. 44, 559–570; 10.1002/hbm.26084 (2023).

52. Desmurget, M. & Sirigu, A. Conscious motor intention emerges in the inferior parietal lobule. Curr. Opin. Neurobiol. 22, 1004–1011; 10.1016/j.conb.2012.06.006 (2012).

53. Ortuño, F. et al. Sustained attention in a counting task: Normal performance and functional neuroanatomy. NeuroImage 17, 411–420; 10.1006/nimg.2002.1168 (2002).

54. Downing, P. E., Jiang, Y., Shuman, M. & Kanwisher, N. A cortical area selective for visual processing of the human body. Science 293, 2470–2473; 10.1126/science.1063414 (2001).

55. Schwarzlose, R. F., Baker, C. I. & Kanwisher, N. Separate face and body selectivity on the fusiform gyrus. J. Neurosci. 25, 11055–11059;10.1523/JNEUROSCI.2621-05.2005 (2005).

56. Peelen, M. V. & Downing, P. E. The neural basis of visual body perception. Nat. Rev. Neurosci. 8, 636–648; 10.1038/nrn2195 (2007).

57. Vocks, S. et al. Differential neuronal responses to the self and others in the extrastriate body area and the fusiform body area. Cogn. Affect. Behav. Neurosci. 10, 422–429; 10.3758/CABN.10.3.422 (2010).

58. Eickhoff, S. B. et al. Anatomical and functional connectivity of cytoarchitectonic areas within the human parietal operculum. J. Neurosci. 30, 6409–6421; 10.1523/JNEUROSCI.5664-09.2010 (2010).

59. Del Vecchio, M. et al. Tonic somatosensory responses and deficits of tactile awareness converge in the parietal operculum. Brain 144, 3779–3787; 10.1093/brain/awab384 (2021).

60. Sirigu, A. & Desmurget, M. Somatosensory awareness in the parietal operculum. Brain 144, 3558–3560; 10.1093/brain/awab415 (2021).

61. Naito, E., Ehrsson, H. H., Geyer, S., Zilles, K. & Roland, P. E. Illusory arm movements activate cortical motor areas: A positron emission tomography study. J. Neurosci. 19, 6134–6144; 10.1523/JNEUROSCI.19-14-06134.1999 (1999).

62. Annett, J. Motor imagery: Perception or action? Neuropsychologia 33, 1395–1417; 10.1016/0028-3932(95)00072-b (1995).

63. Annett, J. On knowing how to do things: a theory of motor imagery. Brain Res. Cogn. Brain Res. 3, 65–69; 10.1016/0926-6410(95)00030-5 (1996).

64. Grush, R. The emulation theory of representation: Motor control, imagery, and perception. Behav. Brain Sci. 27, 377–396; discussion 396-442;10.1017/s0140525x04000093 (2004).

65. Scheperjans, F., Palomero-Gallagher, N., Grefkes, C., Schleicher, A. & Zilles, K. Transmitter receptors reveal segregation of cortical areas in the human superior parietal cortex: Relations to visual and somatosensory regions. Neuroimage 28, 362–379; 10.1016/j.neuroimage.2005.06.028 (2005).

66. Scheperjans, F. et al. Observer-independent cytoarchitectonic mapping of the human superior parietal cortex. Cereb. Cortex 18, 846–867; 10.1093/cercor/bhm116 (2008).

67. Fadiga, L. et al. Corticospinal excitability is specifically modulated by motor imagery: A magnetic stimulation study. Neuropsychologia 37, 147–158; 10.1016/s0028- 3932(98)00089-x (1999).

68. Kimura, N., Furuta, T., Miura, G. & Naito, E. Combining motor imagery and action observation with vibratory stimulation increases corticomotor excitability in healthy young adults. J. Behav. Brain Sci. 12, 177–195; 10.4236/jbbs.2022.125010 (2022).

69. Arroyo, S. et al. Functional significance of the mu rhythm of human cortex: An electrophysiologic study with subdural electrodes. Electroencephalogr. Clin. Neurophysiol. 87, 76–87; 10.1016/0013-4694(93)90114-b (1993).

70. Duann, J. R. & Chiou, J. C. A comparison of independent event-related desynchronization responses in motor-related brain areas to movement execution, movement imagery, and movement observation. PLOS ONE 11, e0162546; 10.1371/journal.pone.0162546 (2016).

71. Pascual-Leone, A. et al. Modulation of muscle responses evoked by transcranial magnetic stimulation during the acquisition of new fine motor skills. J. Neurophysiol. 74, 1037–1045; 10.1152/jn.1995.74.3.1037 (1995).

72. Karni, A. et al. Functional MRI evidence for adult motor cortex plasticity during motor skill learning. Nature 377, 155–158; 10.1038/377155a0 (1995).

73. Debarnot, U., Clerget, E. & Olivier, E. Role of the primary motor cortex in the early boost in performance following mental imagery training. PLOS ONE 6, e26717; 10.1371/journal.pone.0026717 (2011).

74. Makary, M. M., Eun, S. & Park, K. Greater corticostriatal activation associated with facial motor imagery compared with motor execution: A functional MRI study. NeuroReport 28, 610–617; 10.1097/WNR.0000000000000809 (2017).

75. Oldfield, R. C. The assessment and analysis of handedness: The Edinburgh inventory. Neuropsychologia 9, 97–113; 10.1016/0028-3932(71)90067-4 (1971).

76. Ashburner, J. VBM tutorial, (2010).

77. Morita, T. et al. Hyper-adaptation in the human brain: Functional and structural changes in the foot section of the primary motor cortex in a top wheelchair racing Paralympian. Front. Syst. Neurosci. 16, 780652; 10.3389/fnsys.2022.780652 (2022).

78. Hu, X. et al. Voxel-based morphometry studies of personality: Issue of statistical model specification--effect of nuisance covariates. Neuroimage 54, 1994–2005; 10.1016/j.neuroimage.2010.10.024 (2011).

79. Ridgway, G. R. et al. Issues with threshold masking in voxel-based morphometry of atrophied brains. Neuroimage 44, 99–111; 10.1016/j.neuroimage.2008.08.045 (2009).

80. Power, J. D., Barnes, K. A., Snyder, A. Z., Schlaggar, B. L. & Petersen, S. E. Spurious but systematic correlations in functional connectivity MRI networks arise from subject motion. Neuroimage 59, 2142–2154; 10.1016/j.neuroimage.2011.10.018 (2012).

81. Evans, A., Kamber, M., Collins, D. & MacDonald, D. An MRI-based probabilistic atlas of neuroanatomy in Magnetic resonance scanning and epilepsy 263-274 (Plenum Press, 1994).

82. Friston, K. J. et al. Analysis of fMRI time-series revisited. Neuroimage 2, 45–53; 10.1006/nimg.1995.1007 (1995).

83. Worsley, K. J. & Friston, K. J. Analysis of fMRI time-series revisited--again. Neuroimage 2, 173–181; 10.1006/nimg.1995.1023 (1995).

84. Holmes, A. P. & Friston, K. J. Generalisability, random effects and population inference. in vol. NeuroImage 7, S754; 10.1016/S1053-8119(18)31587-8 (1998).

85. Eickhoff, S. B. et al. A new SPM toolbox for combining probabilistic cytoarchitectonic maps and functional imaging data. Neuroimage 25, 1325–1335; 10.1016/j.neuroimage.2004.12.034 (2005).

86. Amunts, K., Mohlberg, H., Bludau, S. & Zilles, K. Julich-Brain: A 3D probabilistic atlas of the human brain’s cytoarchitecture. Science 369, 988–992; 10.1126/science.abb4588 (2020).

87. McLaren, D. G., Ries, M. L., Xu, G. & Johnson, S. C. A generalized form of context-dependent psychophysiological interactions (gPPI): A comparison to standard approaches. Neuroimage 61, 1277–1286; 10.1016/j.neuroimage.2012.03.068 (2012).

88. Whitfield-Gabrieli, S. & Nieto-Castanon, A. A. Conn: A functional connectivity toolbox for correlated and anticorrelated brain networks. Brain Connect. 2, 125–141; 10.1089/brain.2012.0073 (2012).

89. Behzadi, Y., Restom, K., Liau, J. & Liu, T. T. A component based noise correction method (CompCor) for BOLD and perfusion based fMRI. NeuroImage 37, 90–101; 10.1016/j.neuroimage.2007.04.042 (2007).

